# Loop detection using Hi-C data with HiCExplorer

**DOI:** 10.1101/2020.03.05.979096

**Authors:** Joachim Wolff, Rolf Backofen, Björn Grüning

## Abstract

Chromatin loops are an important factor in the structural organization of the genome. The detection of chromatin loops in Hi-C interaction matrices is a challenging and compute intensive task. The presented approach shows a chromatin loop detection algorithm which applies a strict candidate selection based on continuous negative binomial distributions and performs a Wilcoxon rank-sum test to detect enriched Hi-C interactions.

## Introduction

Chromosome conformation capture (3C) (1) and its successors 4C (2, 3), 5C (4) and Hi-C (5) are protocols to study the three dimensional structure of a genome. With Hi-C data a genome wide interaction map of the chromatin can be created and chromatin loops can be inferred. Chromatin loops reflect the interaction of promoters and enhancers, gene loops, architectural loops or polycomb-mediated regions (6) and can be detected as enriched regions in comparison to their neighborhood. By identifying these regions it can be shown that there are long range regulations that impacts e.g. the directionality of RNA synthesis (7) or the long distance between cis-regulatory elements (6) that can not be explained and shown otherwise. Based on Rao (8) the genomic distance between two loci is usually limited to ~ 2 megabases (Mb).

Different algorithms are able to detect loops: HiCCUPS uses a *donut algorithm*; which considers all elements of a Hi-C interaction matrix as peaks and tests if the region around them is significantly different from the neighboring interactions. HiCCUPS is part of the software Juicer (https://github.com/aidenlab/juicer), and the implementation requires a general purpose GPU (GPGPU), which imposes a barrier to many users by simply not having access to an Nvidia GPU. However, an experimental CPU based implementation was released too. HOMER (9) creates a relative contact matrix per chromosome and scans these for locally dense regions. HOMER does not support standard file formats for Hi-C matrices like *cool*, which imposes the need to create all data from scratch which is time consuming and is a potential source of errors and inaccuracies. GOTHIC (10) models the probability of two genomic locations to interact with each other as a mix of different biases and the chance of random interactions. The problem is that GOTHIC detects a large number of significant interactions but is not able to detect only the enriched regions in relation to their neighborhood. It is a good tool to detect significant interactions in a Hi-C interaction matrix but it is not suitable for the specific task of chromatin loop detection. cLoops (11) uses a DBSCAN based approach in combination with a local background to estimate the statistical significance of a loop. cLoops is mainly designed for HiChIP data and not for Hi-C. With HiChIP protein binding sites can be investigated in their 3D context, however, similar with promoter capture Hi-C only the targeted regions are enriched. The consequence of this is an Hi-C matrix with data only available at these enriched regions and foreknowledge of potential loop locations is required. FastHiC (12) is a loop detection algorithm based on a hidden Markov random field Bayesian from (13), which focuses on intra topological associated domain (TAD) loops in a range of 40kb and therefore not on chromatin loops outside of TADs.

Here we present an algorithm that is able to detect Hi-C loops. It is optimized for a high parallelization by providing the option to assign one thread per chromosome and multiple threads within a chromosome. This approach makes fully use of the resources available in the last generation of multi-core CPU platforms.

## Methods

According to Rao (8) the majority of the anchor points of detected loops lies within a range of 2Mb. This insight can be used to decrease the search space in a biologically meaningful way and also reduces the computational burden, maintaining a low memory footprint at the same time. Moreover, interaction pairs with genomic distances which are too close to each other and therefore quite close to the main diagonal have already high interaction counts. It is in many cases unlikely that these pairs contribute enrichments in the context of their neighborhood. This observation can be explained by the high interaction count between two loci, the closer they are in the one dimensional space and are therefore close to the main diagonal. To detect intra-TAD enrichments specialized algorithms like FastHiC should be used. A general problem for Hi-C interactions with few absolute counts is the difficulty to determine if their interactions are true interactions or noise. These artifacts cannot be corrected by the used Hi-C interaction matrix correction algorithms like iterative correction and eigenvector decomposition (ICE) (14) or Knight-Ruiz (KR) (15). These algorithms perform a matrix balancing and correct for an uneven distribution of the interaction counts per genomic position. The correction algorithms are not able to decide, and therefore filter out, if interactions are true interactions or noise. To account for these known problems in the Hi-C interaction data all values below a given threshold are discarded and noise is removed.

### A. Algorithm

A strict candidate selection is key to reduce the computational complexity for the loop detection algorithm. To take the observation from Rao into account, a minimum and maximum loop size can be defined to restrict the search space (Figure 1B). In Hi-C the main data structure is the symmetrical *n* × *n* interaction count matrix (ICM):

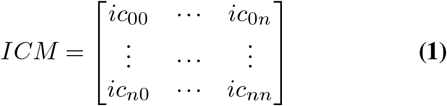

The relative genomic distance is given by:

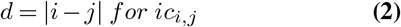

And *ic_i,j_* as an element of Hi-C interaction matrix *ICM*.

**Fig. 1.**
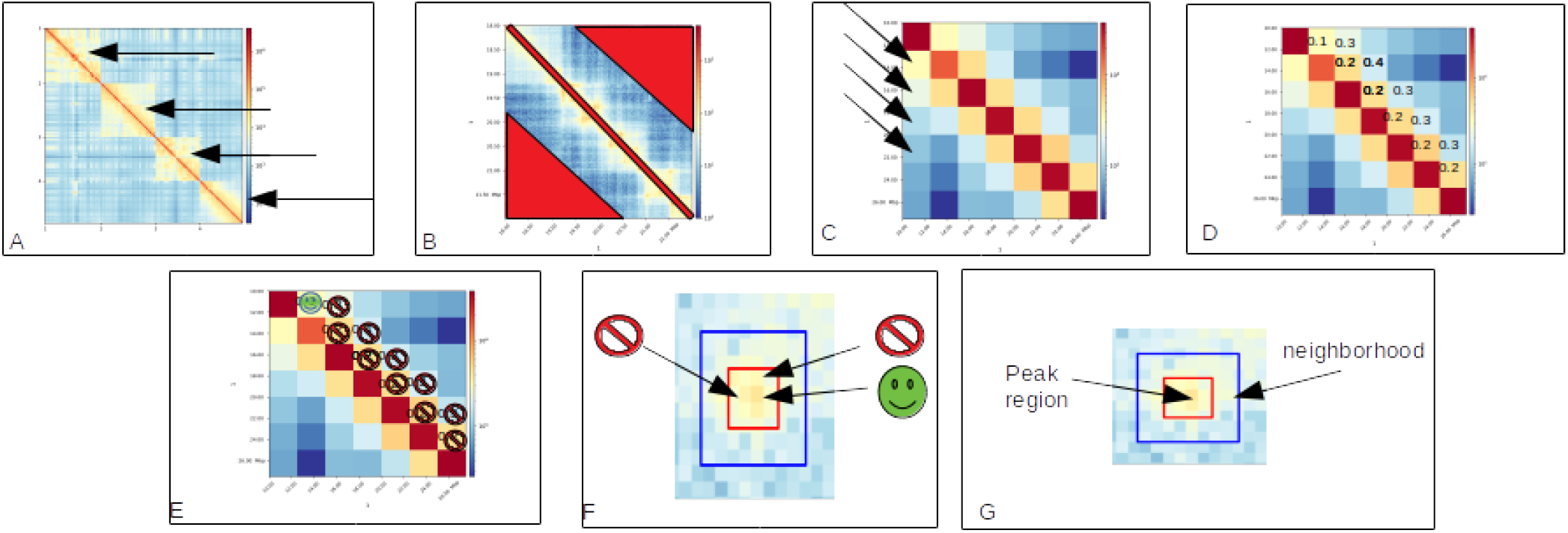
Graphical representation of the loop detection algorithm: A: Compute each chromosome independently. B: Accept an interaction if their relative distance is within: *minLoopSize < relative distance < maxLoopSize*. C: Fit cNB distribution per relative distance. D: Compute a p-value for each interaction. E: Reject candidate if the p-value is too high or the interaction value is smaller a) a fixed threshold or b) a fixed percentage of the maximum value of its relative distance. F:Define neighborhood around an interaction. Accept as candidate the one with the highest interaction. G: Smooth neighborhood and peak region. Reject candidate if: a) maximum of peak region is smaller than the maximum of the neighborhood, b) the mean of the peak region is smaller than the mean of the neighborhood or c) the p-value computed by Wilcoxon rank-sum test comparing peak and neighborhood region is too high.

**Fig. 2.**
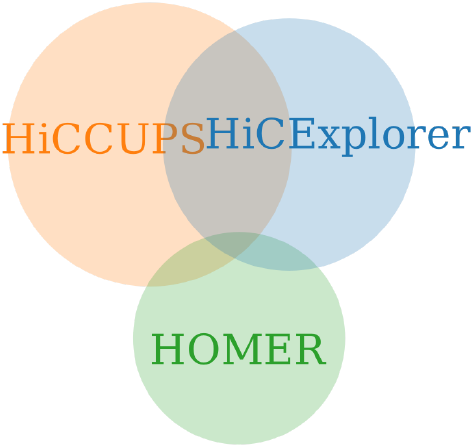
Detected loops by different software.

**Fig. 3.**
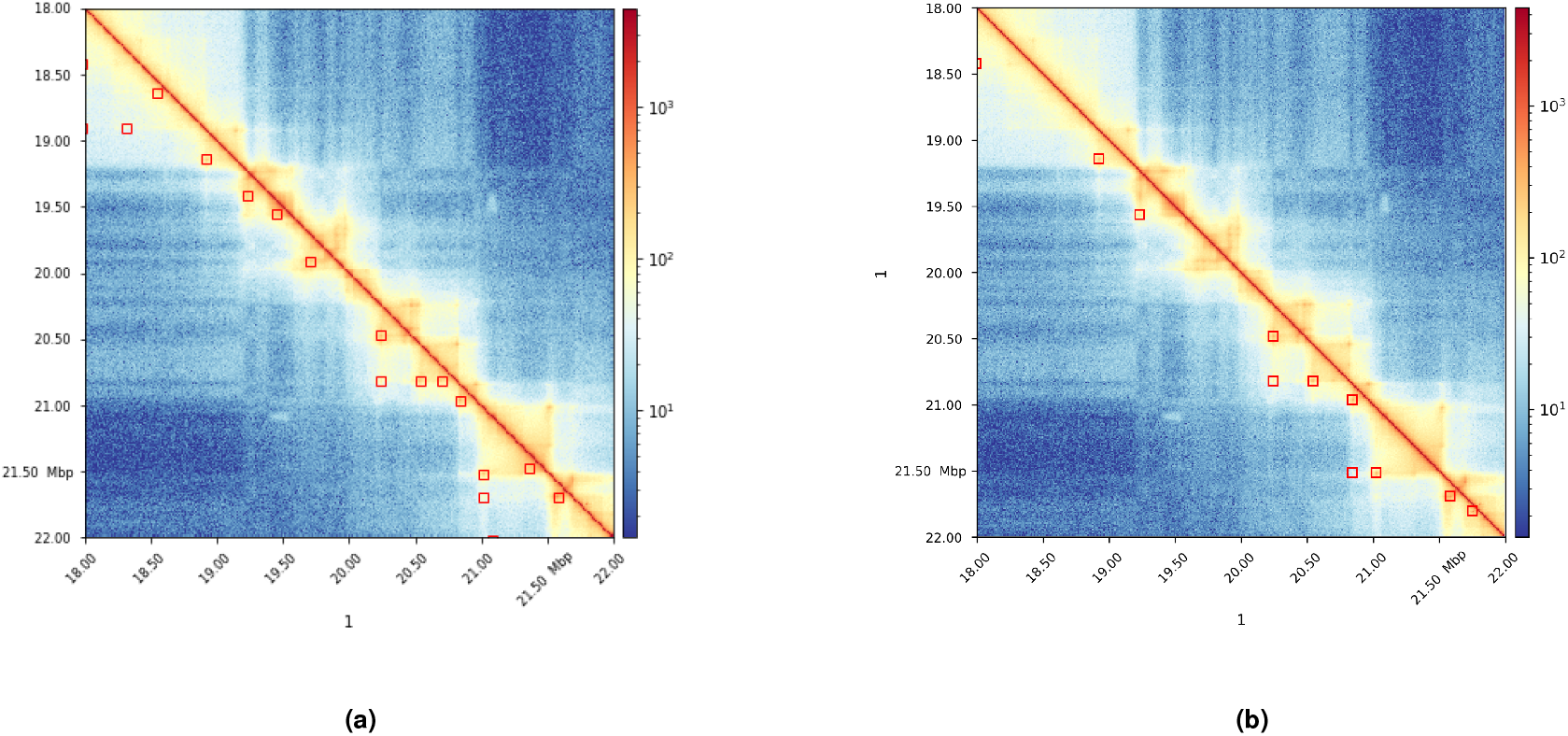
Plot of chr1 18 - 22 MB on GM12878. **(a)**: Detect loops from HiCExplorer, **(b)**: HiCCUPS. Plot with HiCExplorer hicPlotMatrix.

#### A.1. Candidate selection per genomic distance

To detect enriched Hi-C interactions the Hi-C data is fitted per genomic distance d independently to a continuous negative binomial distribution, (Figure 1C):

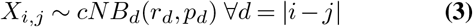

To make the continuous negative binomial function continuous the binomial coefficient must be replaced as it used by edgeR (16, 17) and was discussed at stackoverflow^1^:

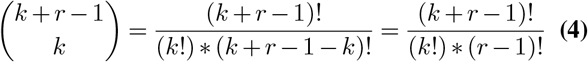

The gamma function is defined for any 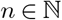:

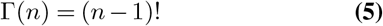

Moreover, the gamma function is defined for any *n* ∈ ℝ_>0_:

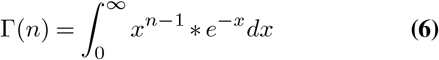

With Equation (5) the binomial coefficient can be reformulated as:

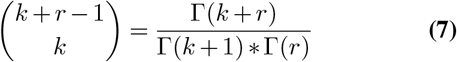

Which leads to the probability mass function for a ‘continuous negative binomial distribution’ with ∀*k* ∈ ℝ_>0_ and ∀*r* ∈ ℝ_>0_:

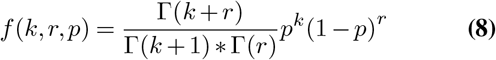

The probability of observing an interaction count or a higher one at the genomic distance d is given by the continuous negative binomial cumulative density:

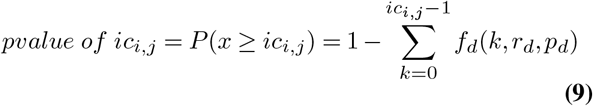

Only the interaction counts with p-values smaller than a threshold are accepted as candidates (Figure 1D and 1E); these candidates are further filtered to remove candidates with too few absolute interactions (Figure 1E). To account for the decreasing contact probabilities for an increasing genomic distance, a distance dependent threshold is used. The maximum interaction count per genomic distance is considered and each candidate is removed if it has less interactions as the maximum interaction count times a threshold percentage, e.g. all interactions are removed if they have less interactions counts as 10% of the maximum value of their genomic distance.

#### A.2. Loop peak detection

To detect enriched regions in a Hi-C interaction matrix the entire neighborhood needs to be considered. A neighborhood is a square of size n with the candidate element in its center, see Figure 1F. An enriched region needs to have an enriched interaction count in relation to the elements in its neighborhood. The concept of a neighborhood comes with a few issues: First, in one neighborhood there can be multiple candidates detected from different, but next to each other located genomic distances. Second, if a candidate is significant for its genomic distance it is not necessarily an enriched value for its neighborhood. And third, a single enriched interaction in a neighborhood is possible, but is likely a false positive. Meaningful enriched interactions appear in groups and form a peak in the two dimensional space, as shown in Figure 1F. To handle the first issue, all candidates in one neighborhood are pooled together, only the candidate with the highest interaction count for one neighborhood is considered to be a representative of its neighborhood; all others are dropped (Figure 1F). To cover the second and third issue, the remaining candidate neighborhoods are smoothed using a sliding window approach in x and y directions to remove possible outliers. Moreover, the neighborhood is split into a peak and a background region by considering the square around the candidate as the peak region and the remaining elements of the neighborhood as the background, see Figure 1G. The peak and neighborhood square sizes are defined by their inradius values, *peakWidth* and *windowSize*. All candidates which fulfill either of the following two conditions are rejected: a) *mean*(*background*) ≥ *mean*(*peak*) or b) *max* ≥ (*background*) ≥ *max*(*peak*). This filter step is necessary to address the candidate peak value is a singular outlier within the neighborhood and after smoothing there is no more peak. The last case can occur for candidates which had no overlapping neighborhoods of candidates and the highest interaction count in their neighborhood was never considered as a significant outlier by its continuous negative binomial distribution. Furthermore, the Wilcoxon rank-sum test with H0 hypothesis background and peak regions are from the same distribution with significance level p is used. The mentioned filter steps guarantee only neighborhoods with a centering peak value are considered.

## Results

To verify the results of the chromatin loop detection algorithm it was tested on Hi-C data on various cell types published by Rao 2014: GM12878, K562, IMR90, HUVEC, KBM7, NHEK and HMEC. Additionally, the detected chromatin loop locations are correlated with binned protein peak locations of the 11-zinc finger protein CTCF. CTCF is a known loop binding factor (8) although not all peaks need to have CTCF attached (18), especially in the case of a gene or a polycomb-mediated loop (6). An overlap of a detected chromatin loop region was accepted if at both loci CTCF was detected. CTCF was matched to the GM12878, HMEC, HUVEC,K562 and NHEK cell sample, for IMR90 and KBM7 no CTCF from the same source is provided. HiCExplorer’s implementation is tested against HiCCUPS algorithm from the Juicer software and HOMER’s loop detection. The algorithms of GOTHIC, cLoops, FastHiC are not part of the comparison due to different focuses of the algorithms.

### B. HiCExplorer candidate selection

In the following section, the results were computed on GM12878 with applied Knights-Ruiz correction and a 10kb fixed bin size resolution. The loop detection considers each chromosome independently, we will use data from chromosome 1 to show the search space reduction as an example. The p-value was set to 0.05 for continuous negative binomial distribution candidate selection, a minimal interaction peak height of 20, a peak width of 6 and a window size of 10 and a maximal interaction count share of 0.1. Based on the Rao’s observation that the maximum distance of two loci forming a loop usually does not exceed 2Mb, the upper boundary was set to this value. The lower limit for genomic distances of two loci was set to 100kb. The upper and lower distance settings decrease the search space from 40.5 million to 3.9 million candidates. The million candidates are given by the count of non-zero interactions. However, the parameters for minimum and maximum distance between two loci are adjustable. The p-value selection based on continuous negative binomial distributions with level 0.05 reduces the search space from 3.9 million to 530,000 candidates for chromosome 1. A pruning of the candidates with less absolute interactions than maximum interaction count share of 0.1 further decreases the search space to 82,000 candidates. The candidate pooling per neighborhood decreases the search space again to only 3515 candidates and gives a vastly small number to apply the testing with Wilcoxon rank-sum test. This shows that a good candidate selection helps to decrease the search space drastically. Starting from 40.5 million candidates the Wilcoxon rank-sum test gets only 0.00008% of the original candidates to test.

For other cell lines published by Rao 2014 the situation is comparable (Table 5). For all cell lines the number of detected candidates is of the same order of magnitude, which indicates a robust candidate selection with the chosen continuous negative binomial distributions. Another important aspect to reduce the search space is the observation that peaks in Hi-C interaction matrices have a two dimensional area and not single elements. Peaks are only detectable in the context of their local neighborhood as the significance given by the continuous negative binomial distributions is not enough. This leads to multiple candidates per neighborhood and consequently, only the one with the highest interaction count can be considered as the peak. The pooling of the candidates under these conditions leads to a reduction of the search space in GM12878 cells of a factor of 23. The reduction rates on the other cell types are similar. However, the situation is different after the testing of the peak region (Table 1). The number of detected loops differs between 3000 to 10,000 loops. To explain this different detection behaviour the non-zero values and implicitly the read coverage per bin is considered. This shows the relation, the higher the read coverage, the more regions are detected (see Table 1 and 5). However, the comparison of IMR90 and KBM7 shows that the level of matrix sparsity is not the only explanation, the samples are from different cell types and therefore different structures can be formed. The read coverage effect can be explained by the preprocessing of the neighborhood region before it is separated into a peak and background regions: The neighborhood is smoothed in x and y direction; therefore the lower the read coverage, the more likely it is to have ‘holes’ i.e. elements with zero interactions in a neighborhood and with this, the neighborhood is smoothed more into the direction of the zero values and become overall more equal. To have close to equal distributed values is contrary to a peak, therefore more candidates are rejected by the statistical test. The candidate selection approach via the definition of a neighborhood makes the algorithm sensitive to the resolution of the Hi-C interaction matrix. The lower the resolution, the smaller the neighborhood needs to be. Otherwise the chances of having elements in the neighborhood which are peaks or TADs, or even the main diagonal are too high. Decreasing the size of the neighborhood creates at the same time another issue: the neighborhood and therefore the number of elements in the peak and background regions are becoming too less. This leads to non-significant test results, and leads to the insight that first, the neighborhood size needs to be adjusted to the bin resolution of the Hi-C matrix and second, a neighborhood should contain at least around 250 - 300 elements to produce useful results.

**Table 1.**
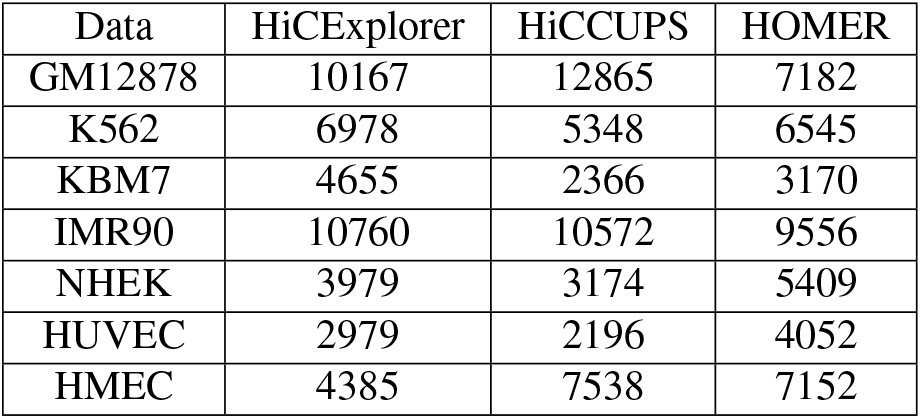
Detected loops on different cell types cells from Rao 2014, with 10kb resolution, HiCExplorer and HiCCUPS with applied KR correction. Used parameters for all three tools are listed in Table 8

**Table 2.** Number of detected loops with CTCF match, percentage in brackets.

| Data | HiCEXplorer | HiCCUPS | HOMER |
| --- | --- | --- | --- |
| GM12878 | 5440 (0.53) | 6354 (0.49) | 1217 (0.16) |
| K562 | 3125 (0.44) | 3219 (0.6) | 1137 (0.17) |
| NHEK | 1544 (0.38) | 1419 (0.44) | 1224 (0.22) |
| HUVEC | 1787 (0.59) | 1260 (0.57) | 864 (0.21) |
| HMEC | 2284 (0.52) | 4323 (0.57) | 1681 (0.23) |

**Table 3.** Number of detected loops with CTCF match, percentage in brackets.

| Data | HiCEXplorer ∩ HiCCUPS | HiCEXplorer ∩ HOMER | HiCCUPS ∩ HOMER | HiCEXplorer ∩ HiCCUPS ∩ HOMER |
| --- | --- | --- | --- | --- |
| GM12878 | 4225 | 490 | 574 | 291 |
| K562 | 2034 | 347 | 298 | 155 |
| KBM7 | 508 | 125 | 68 | 33 |
| IMR90 | 4121 | 931 | 974 | 522 |
| NHEK | 1142 | 333 | 280 | 145 |
| HUVEC | 446 | 140 | 172 | 39 |
| HMEC | 2049 | 436 | 755 | 275 |

**Table 4.** CTCF correlation with intersected loops, percentage in relation to number of intersected loops, see Table 2

| Data | HiCEXplorer ∩ HiCCUPS | HiCEXplorer ∩ HOMER | HiCCUPS ∩ HOMER | HiCEXplorer ∩ HiCCUPS ∩ HOMER |
| --- | --- | --- | --- | --- |
| GM12878 | 3344 (0.79) | 353 (0.72) | 385 (0.67) | 238 (0.81) |
| K562 | 1533 (0.73) | 214 (0.61) | 205 (0.68) | 117 (0.75) |
| NHEK | 709 (0.62) | 192 (0.57) | 162 (0.57) | 91 (0.62) |
| HUVEC | 390 (0.87) | 114 (0.81) | 124 (0.72) | 39 (1.0) |
| HMEC | 1500 (0.73) | 312 (0.71) | 530 (0.70) | 217 (0.78) |

**Table 5.** Initial possible candidates vs. reduced candidate set of HiCExplorer for chromosome 1.

| Data | Initial candidates | Candidates for peak detection |
| --- | --- | --- |
| GM12878 | 61.8 mio | 1722 |
| K562 | 19.2 mio | 2948 |
| KBM7 | 14.2 mio | 2321 |
| IMR90 | 19.3 mio | 2948 |
| NHEK | 10.1 mio | 2384 |
| HUVEC | 7.6 mio | 3249 |

#### B.1. Comparison to HiCCUPS and HOMER

The number of detected enriched regions of HiCExplorer, HiCCUPS and HOMER differs between the samples, the number of detections is on a comparable level (Table 1). The situation is different for the successful overlapped binned peaks with CTCF. HICCUPS is in terms of accuracy the best, for three cell lines (NHEK, HUVEC, HMEC) it is slightly (5 - 7%), for K562 a lot (18%) better; for GM12878 HiCExplorers accuracy is better by 4%. The number of correctly detected loops is usually the highest for HiCCUPS, HiCExplorer is for K562, NHEK, HUVEC on a comparable level or slightly higher. On GM12878 HiCExplorer detects around 900 loops less and on HMEC HiCCUPS detects 2000 more loops. The performance by HOMER in terms of quantity and quality are not comparable with HiCExplorer and HiCCUPS. The accuracy is between 16 - 23 % where HiCExplorer has 38 - 53% and HiCCUPS 44 - 60%. Moreover, the number of correct loops is significantly less: HOMER detects between 864 - 1681 loops, HiCExplorer 1220 - 5440 and HiCCUPS 1266 - 6354 (Table 2). It needs to be mentioned that the correlation with CTCF can only give an indication of the quality of the detected loops. First, loop structures representing gene or polycomb-mediated loops do not have CTCF at their anchor points. Second, the used method with ChIP-Seq data is biased and not available in a two dimensional space. HiChIP data could be used for a better benchmark but was not available.

In comparison to HiCCUPS HiCExplorer misses the 2% chromatin loops stated in Rao 2014 for genomic distances > 2 Mb which should include inter-chromosomal enrichments. These inter-chromosomal enrichments are not detectable by HiCExplorer because each chromosome is computed independently. In our testing also HiCCUPS was not able to detect non inter-chromosomal interaction. Recomputed results on GM12878 with HiCCUPS and three resolutions, 5kb, 10kb and 25kb, in total 17768 loops were detected and 4910 have a distance greater 2 Mb; on 10kb out of 12865 loops, 2968 have a greater distance than 2 Mb. Contrastly, it is not quite clear on which base Rao 2014 states that only 2% of the loops are in 2 Mb range. However, if the correlated loops are computed on HiCCUPS data with all loops of distances greater 2 Mb are removed, 6205 instead of 6354 loops can be correlated with CTCF. This supports the restriction to a range of 2 Mb. If the restriction of the genomic distance between two loci from 100kb to 2Mb is removed for HiCExplorer and all intra-chromosomal contacts are considered, the number of candidates to be tested increases by a factor of 10, and the number of accepted peaks increased from 10167 to 11723. The proposed peak detection algorithm was tested on GM12878 primary dataset with a 10kb resolution and performs on a similar level as Juicers HiCCUPS CPU version, 3:40 minutes runtime in comparison to 3 minutes. However, the memory usage is with 26 GB higher compared to 11 GB. Juicers HiCCUPS GPU version is either significant slower (9 minutes) if it is not restricted in the search distance as HiCCUPS CPU version or, if it is restricted, outperforms both with a runtime of 1:40 minutes.

The runtime of HOMER on the 10kb resolution with more than 8 hours is extensively longer. The memory usage of HiCExplorer and HiCCUPS perform on a similar level, while HOMER demands more than 52GB (see Table 6). HOMER offers the use of a parallelization per chromosome, but the high peak memory usage of 52 GB makes it impractical to use for many users.

**Table 6.** Computed on GM12878 primary with KR on 10kb resolution, on 2xIntel XEON E5-2630 v4 @ 2.20GHz 2×10 cores / 2×20 threads, 120 GB memory with Nvidia Tesla T4. HiCCUPS GPU restricted and CPU version are restricted to 8 MB distance, HiCExplorer was computed its default distance of 2 MB; however HiCExplorer supports other distances too.

| Tool | Runtimes | Memory |
| --- | --- | --- |
| HiCEXplorer (4*4 threads) | 7:57 min | 13 GB |
| HiCEXplorer (16*12 threads) | 3:41 min | 26 GB |
| HiCCUPS GPU | 8:59 min | 11.6 GB |
| HiCCUPS GPU restricted | 1:41 min | 11.3 GB |
| HiCCUPS CPU (40 threads) | 3:03 min | 11.8 GB |
| HiCCUPS CPU (4 threads) | 11:21 min | 9.8 GB |
| HOMER | 8:51 h | 52 GB |

**Table 7.** Sparsity level of the 10 kb Hi-C interaction matrices. The dense matrix contains 309,581 × 309,581 elements.

| Data | Non-zero elements | Sparsity |
| --- | --- | --- |
| GM12878 | 1,810 mio | 0.0189 |
| K562 | 781 mio | 0.0081 |
| KBM7 | 465 mio | 0.0048 |
| IMR90 | 415 mio | 0.0043 |
| NHEK | 348 mio | 0.0036 |
| HUVEC | 268 mio | 0.0028 |
| HMEC | 188 mio | 0.0019 |

The overlap of detected peaks of HiCExplorer, HICCUPS and HOMER is quite different. HiCExplorer and HOMER share a quarter to half of detect loops, while the share with the detected loops with HOMER is 5 - 10%. HICCUPS and HOMER share a different loops but on a same level as HiCExplorer and HOMER. The intersection of all three is around half of HiCExplorer and HOMER or HICCUPS and HOMER (Table 1 and 3). The correlated CTCF for all intersected results is with 60 - 82% higher as with the individual software alone (Table 2 and 4).

## Discussion

The search space of an algorithm is the dominating factor for its accuracy and performance. Therefore, pruning it should be the main goal of newly designed algorithms. Bruteforce solutions like HiCCUPS with no restrictions to the search space are, in theory, able to detect all possible enriched regions, but at a cost of a hardware demanding implementation. HiCCUPS solved this by the massive parallel computational resources via GPGPU. The limitation of the search space to a genomic distance of 100kb to 2 Mb has only a small impact to the detected peaks. HOMER, however, has limitations on the search space but detects less number of loops and the detected ones have a significantly lower correlation over all samples to CTCF. Moreover, HOMER does support a parallelization per chromosome like HiCExplorer but is significantly slower than HiCExplorer by using extensively more memory per core. Furthermore, it could be shown that the sparsity and therefore read coverage of a Hi-C interaction matrix has a major influence on the detection of peaks in their neighborhood. The sparser a Hi-C interaction matrix is, the more likely it is that possible valid regions detected by the continuous negative binomial distribution filtering are rejected by Wilcoxon rank-sum test.

## Acknowledgements

We thank Simon Bray and Anup Kumar for proof reading the manuscript.

## Availability of data and materials

Hi-C data: GSE63525; Rao 2014. ChIP-Seq data: CTCF from Bernstein (GSM733752) The implementation of this algorithm is part of the software HiCExplorer (19, 20) since version 3.2. HiCExplorer is licenced under GPLv3 and is available on Github (https://github.com/deeptools/HiCExplorer/) or as conda package in the bioconda channel (21). HiCExplorer is implemented in Python 3.

## Supplementary Note 1: Used parameters for results

**Table 8.** Setting to compute loops with HiCExplorer, HiCCUPS and HOMER.

| Data | HiCEXplorer | HiCCUPS | HOMER |
| --- | --- | --- | --- |
| GM12878 KR primary<br>+<br>replicate | -pit 20 -windowSize 8 -pp 0.01 -p 0.01<br>-t 16 -tpc 12 -minLoopDistance 100000 -mip 0.1<br>-st wilcoxon-rank-sum<br>-o gm12878_10kb.bedgraph | -r 10000 -k KR | findTADsAndLoops.pl<br>find gm12878_tag/ -cpu 5<br>-res 10000 -window 15000<br>-genome hg19 |
| HMEC | -p 1e-3 -pp 0.01<br>-o hmec_10kb_loops.bedgraph<br>-t 16 -tpc 12 -st wilcoxon-rank-sum -pit 10 -mip 0.01<br>-windowSize 8 | -r 10000 -k KR | findTADsAndLoops.pl<br>find hmec_tag/ -cpu 5<br>-res 10000 -window 15000<br>-genome hg19 |
| HUVEC | -p 1e-3 -pp 0.001<br>-o huvec_10kb_loops.bedgraph<br>-t 16 -tpc 12 -st wilcoxon-rank-sum -pit 10 -mip 0.1<br>-windowSize 8 | -r 10000 -k KR | findTADsAndLoops.pl<br>find huvec_tag/ -cpu 5<br>-res 10000 -window 15000<br>-genome hg19 |
| IMR90 | -pit 7 -windowSize 8 -pp 0.025<br>-p 0.0001 -o imr90_10kb_test.bedgraph -t 16 -tpc 12<br>-minLoopDistance 100000 -mip 0.1<br>-st wilcoxon-rank-sum | -r 10000 -k KR | findTADsAndLoops.pl<br>find imr90_tag/ -cpu 5<br>-res 10000 -window 15000<br>-genome hg19 |
| K562 | -pit 10 -windowSize 8<br>-pp 0.01 -p 0.01<br>-o k562.bedgraph -t 16 -tpc 12<br>-minLoopDistance 100000 -mip 0.1<br>-st wilcoxon-rank-sum | -r 10000 -k KR | findTADsAndLoops.pl<br>find k562_tag/ -cpu 5<br>-res 10000<br>-window 15000<br>-genome hg19 |
| KBM7 | -pit 10 -windowSize 8<br>-pp 0.01 -p 0.01<br>-o kbm7_10kb.bedgraph -t 16 -tpc 12<br>-minLoopDistance 100000 -mip 0.1<br>-st wilcoxon-rank-sum | -r 10000 -k KR | findTADsAndLoops.pl<br>find kbm7_tag/ -cpu 5<br>-res 10000 -window 15000<br>-genome hg19 |
| NHEK | -p 1e-3 -pp 0.0025 -t 16 -tpc 12<br>-st wilcoxon-rank-sum -pit 10<br>-mip 0.1 -windowSize 8 | -r 10000 -k KR | findTADsAndLoops.pl<br>find nhek_tag/ -cpu 5<br>-res 10000 -window 15000<br>-genome hg19 |

## Supplementary Note 2: Software versions

The used HiCExplorer version in this paper is 3.4.2, the used Juicer HiCCUPS version 1.19.02, CUDA 10. HOMER was used in version 4.11.

The data provided by Rao (8) is available as raw data (FASTQ files) or in Juicer’s *hic* format. To process the data with HiCExplorer, the provided *hic* files have been converted to the *cool* format with *hic2cool*^2^. To be able to process the data with HOMER, the raw FASTQ files were used and processed as described in HOMERs documentation.

1 https://stats.stackexchange.com/questions/310676/continuous-generalization-of-the-negative-binomial-distribution/311927

2 https://github.com/4dn-dcic/hic2cool

